# Role of miR-181c in Diet-Induced Obesity Through Regulation of Lipid Synthesis in Liver

**DOI:** 10.1101/2021.08.20.457144

**Authors:** Kei Akiyoshi, Gretha J. Boersma, Miranda D. Johnson, Fernanda Carrizo Velasquez, Brittany Dunkerly-Eyring, Shannon O’Brien, Atsushi Yamaguchi, Charles Steenbergen, Kellie L. K. Tamashiro, Samarjit Das

## Abstract

We recently identified a nuclear-encoded miRNA (miR-181c) in cardiomyocytes that can translocate into mitochondria to regulate mitochondrial gene (mt-COX1), and influence obesity-induced cardiac dysfunction through mitochondrial pathway. Liver plays a pivotal role during obesity. Therefore, we hypothesized that miR-181c plays an important role in pathophysiological complications associated with obesity. We used miR-181c/d^-/-^ mice to study the miR-181c role in lipogenesis in hepatocytes during diet-induced obesity (DIO). Indirect calorimetric measurements were made during the 26 weeks high fat diet (HFD) exposure. qPCR was performed to examine the gene expression involved in lipid synthesis. Here, we show that miR-181c/d^-/-^ mice are not protected against all metabolic consequences of HFD exposure. After 26 weeks of the HFD, miR-181c/d^-/-^ mice had a significantly higher body fat (%) compared to WT. Glucose tolerance tests showed hyperinsulinemia and hyperglycemia, indicative of insulin insensitivity in the miR-181c/d^-/-^ mice. HFD-fed miR-181c/d^-/-^ mice had higher serum and liver triglyceride levels compared to HFD-fed WT mice. qPCR data demonstrated that several genes which are regulated by isocitrate dehydrogenase 1 (IDH1) were upregulated in miR-181c/d^-/-^ liver compared to WT liver. Furthermore, an AAV-8 was used to deliver miR-181c, *in vivo*, to validate the potential role of miR-181c in the liver. miR-181c delivery attenuate the lipogenesis by downregulating the same lipid synthesis genes in the liver. In hepatocytes, miR-181c regulates lipid biosynthesis by targeting IDH1. Taken together, the data indicate that overexpression of miR-181c can be beneficial for various lipid metabolism disorders.

## Introduction

microRNA-181c (miR-181c) is derived from the nuclear genome, translocates to the mitochondria, and more importantly, regulates mitochondrial gene expression and alters mitochondrial function [1–3]. Overexpression of miR-181c in rats causes oxidative stress, which leads to cardiac dysfunction [1], similar to the reported upregulation of miR-181c in human cardiac dysfunction [4]. Recently, it has been identified that myocardial lipid accumulation due to high fat diet (HFD) consumption upregulates miR-181c expression in the heart [5]. Mitochondria activate reactive oxygen species (ROS) production in a type 2 diabetes (T2D) diet-induced obesity (DIO) model, indicating a significant role for mitochondrial dysfunction in glucose homeostasis and insulin resistance [6–8]. As mitochondrial function changes, multiple cellular and physiological functions are affected, contributing to the development of common T2D-related complications.

One miRNA can have different targets in different organs [9, 10]. miR-181c is expressed mainly in the mitochondrial fraction in the heart, and targets a mitochondrial gene (cytochrome c oxidase subunit I, mt-COX1) [1–3]. However, it is possible that miR181c does not localize to the mitochondria in other organs. In fact, it has been shown that miR-181c target genes in a gastric cancer cell line, suggesting that miR-181c mainly localizes to the cytosolic compartment in the gastrointestinal tract [11]. Similarly, miR-181c was found to interact with TNFα in hematopoietic progenitor cells, further suggesting non-mitochondrial localization [12]. It has been demonstrated that miR-181c plays an important role in the priming phase of liver regeneration [13]. miR-181c overexpression has been shown to inhibit Hepatitis C virus replication by directly targeting the 3’-UTR of homeobox A1 (HOXA1) [14].

Normal homeostasis requires a balance between lipid synthesis and lipid oxidation to prevent lipid deposition. In obesity and T2D, lipids, such as triglycerides and total cholesterol, are elevated [15–18]. The role of miRNAs in lipid metabolism is well documented [19]. The liver-specific miRNA, miR-122, has been shown to regulate lipid synthesis by targeting multiple mRNAs that are responsible for lipid biosynthesis, such as FAS, ACC1 and ACC2 [20–22]. SREBF1 regulates cholesterol homeostasis by down-regulating ABC transporters [23]. miR-33 has been shown to alter cholesterol and HDL generation by targeting SREBF1 mRNA [24]. Mitochondrial function directly influences lipid biosynthesis [25–27].

The aim of this study was to identify the role of miR-181c in liver lipid metabolism in the context of obesity. We took advantage of our global miR-181c/d^-/-^ mouse model, which allowed us to investigate liver lipid metabolism in response to the lack of miR-181c during DIO.

## Material and methods

### Animals

Male wild-type (WT) C57/BL6J (Jackson Laboratories, Bar Harbor, ME) and miR-181c/d^-/-^ (c/d KO) mice previously described [3] were used where indicated. Mice were provided *ad libitum* access to standard laboratory chow (CH; 2018 Teklad, Envigo, Frederick, MD) or purified high-fat diet (HF; 60% fat, D12492, Research Diets, New Brunswick, NJ) and tap water. The mice were divided into four experimental groups: WT/CH, WT/HF, c/d KO/CH, and c/d KO/HF (n=6-7 per group). In a subset of high-fat fed mice, a glucose tolerance test was performed at 29 weeks of age. A separate cohort of WT mice were used for the miR-181c overexpression study as described below. All mice were housed in standard polycarbonate cages in a humidity- and temperature-controlled vivarium on a 12 h:12 h light: dark cycle with light onset at 6 am. All procedures were approved by the Institutional Animal Care and Use Committee at the Johns Hopkins University School of Medicine.

### Indirect Calorimetry

Energy expenditure, respiratory exchange ratio, locomotor activity and food intake were determined using indirect calorimetric measurements in an open-flow indirect calorimeter (Oxymax, Columbus Instruments) at 0, 10 and 20 weeks of HF exposure in WT and c/d KO mice. Data were collected for 3 days to confirm acclimation to the calorimetry chambers (stable body weights and food intake), and the third day in the Oxymax was analyzed. Rates of oxygen consumption (VO_2_, ml·kg^−1^·h^−1^) and carbon dioxide production (VCO_2_, ml·kg^−1^·h^−1^) were measured for each chamber every 16 min throughout the study. The respiratory exchange ratio (RER = VCO_2_/VO_2_) was calculated by Oxymax software (v. 4.02) to estimate relative oxidation of carbohydrate (RER = 1.0) vs. fat (RER approaching 0.7), not accounting for protein oxidation.

### IP-Glucose Tolerance Test

Prior to the intraperitoneal-glucose tolerance test (ip-GTT) the mice were fasted overnight (food was removed at 6 pm). Mice were moved into the testing room 1.5 hours prior to glucose injection to habituate (8 am). The ip-GTT was performed according to methods previously described [28]. A baseline blood sample was taken (9:30am) via a small nick of the tail. Hereafter, the mice were injected ip with 1.5 mg/g glucose (20% glucose in sterile water solution) (10 am). Additional blood samples (20 μl) were taken 15, 30, 45, 60, and 120 minutes after glucose injection. Glucose levels in the blood were determined immediately using handheld glucose analyzers (Freestyle; TheraSense, Alameda, CA, USA). Blood was collected using heparinized capillary tubes and stored in micro-centrifuge tubes on ice, centrifuged at 4°C to collect plasma, and plasma was stored at −80°C. Plasma insulin concentration was determined with a commercially available ultra-sensitive mouse insulin ELISA kit (Crystal Chem, Downers Grove, IL, USA).

### Tissue Collection

Animals were deeply anesthetized using isoflurane and blood was collected via cardiac puncture. The heart and liver were rapidly dissected, flash-frozen in liquid nitrogen, and stored at −80 °C for further analysis. Blood was centrifuged at 4°C and plasma was collected and stored at −80 °C.

### Hormone and Triglyceride Analyses

Plasma leptin levels were determined using a commercially available mouse leptin ELISA kit (EMD Millipore, St. Charles, MO, USA). The intra-assay variation of the kit was between 1.06-1.76%, the inter-assay variation of the kit was between 3.01-4.59%. Plasma and liver triglyceride levels were determined using a commercially available colorimetric assay kit (Cayman Chemicals, Ann Arbor, MI, USA). For liver triglyceride levels, samples were further diluted (1:5) and used for triglyceride analysis per the manufacturer’s instructions. Intra- and inter-assay variations of the kit were 1.34% and 3.17%, respectively.

### Mitochondria Isolation Protocol

Freshly isolated mitochondria were prepared from hearts and livers after incubated with RNAlater by differential centrifugation [2]. Briefly, after incubation, the tissues were dissected and placed in Buffer A (in mM: 180 KCl, 2 EGTA, 5 MOPS, 0.2% BSA; pH: 7.25). The tissues were then digested with trypsin (0.0001 g/0.1 g tissue) in 0.7 ml of ice-cold Buffer B (in mM: 225 Mannitol, 75 sucrose, 5 MOPS, 0.5 EGTA, 2 Taurine; pH: 7.25) and finally homogenized in Buffer B with protease inhibitor cocktail (Roche Applied Science, Indianapolis, IN) using a Polytron. To further separate the heart mitochondria from other cellular components and tissue debris, a series of differential centrifugations were performed in a Microfuge 22R centrifuge (Beckman Coulter, Fullerton, CA) at 4°C. The crude pellet was then lysed with QIAzol (Qiagen, Valencia, CA).

### qRT-PCR

Total RNA was isolated from hearts or liver tissues using a miRNeasy kit (Qiagen, Valencia, CA) for miRNA-enriched fraction or RNEasy kit (Qiagen, Valencia, CA) for larger RNA fraction. For the mitochondrial miRNA-enriched fraction a modified protocol was used [2]. In both the cases, a RNase-free DNase kit (Qiagen, Valencia, CA) was used to eliminate genomic DNA contamination. To characterize the integrity of the isolated RNA, spectrophotometric evaluation was performed, using Nanodrop (Thermo Scientific, Wilmington, DE). Only RNA samples with an A260 (absorbance at 260 nm) value greater than 0.15 were used for further experiments. The ratio of the readings at 260 nm and 280 nm (A260/A280) was also measured in order to check the purity of the isolated RNA. For further and more accurate purity and integrity estimation of the isolated RNA, samples were profiled by the Bioanalyzer 2100 (Agilent Technologies, CA). Only RNA with A260/A280 ~2.00 and RIN > 8 was used for the experiments. RNA from heart and liver were reverse transcribed using miScript Reverse Transcription Kit (Qiagen, CA). HiFlex and HiSpec buffers were used for mRNA and miRNA, respectively. PCR for miRNA and mRNA was performed using a miScript SYBR green PCR kit (Qiagen, Valencia, CA) and detected with a CFX96 detector (Bio-Rad, CA). qPCR was performed with primers for miR-181c, SNORD61, and 5S rRNA (Qiagen, CA). All reactions were performed in triplicate. Primer sequences for all other genes are reported in **Table 1**.

**Table 1:**
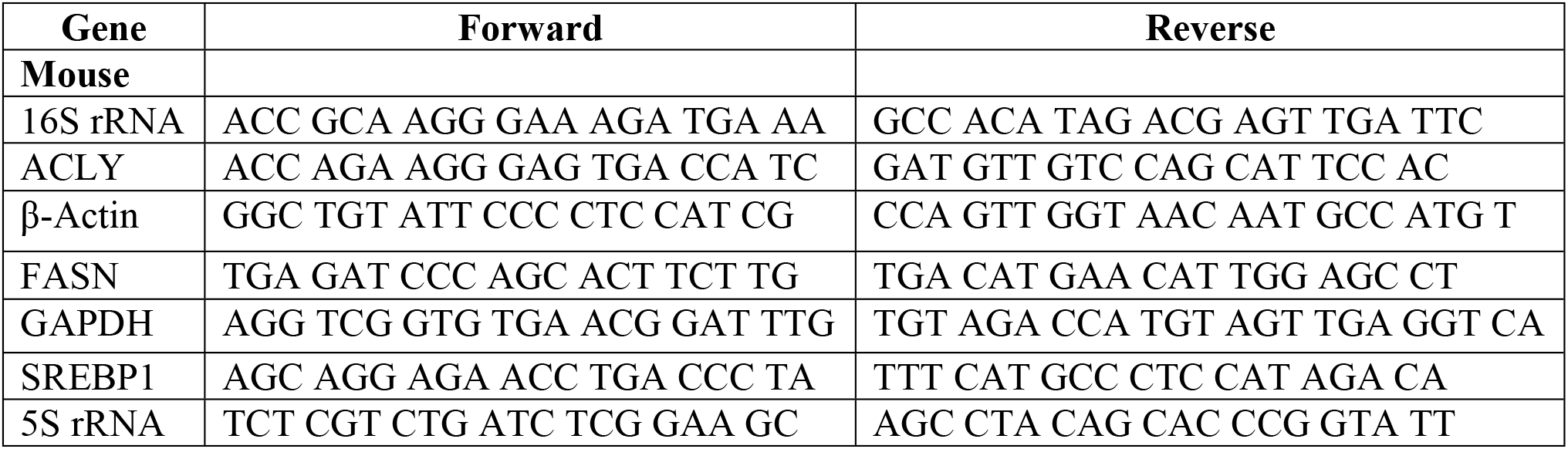
Primers for gene candidates of interest used for qPCR analyses are listed below.

### Western Blot

Liver, heart and isolated mitochondria pellets were lysed with RIPA buffer and protein content was measured by Bradford assay. Cell homogenate protein was separated by 1D gel electrophoresis. After transfer to a PVDF membrane, the membrane was incubated with antibody that recognizes proteins IDH 1 (1:1000) and IDH 2 (1:1000) (Cell Signaling, MA) and α-Tubulin (Abcam, MA) in Tris-Buffered Saline (pH 7.4) with 1% TWEEN 20 (TBS-T) with 5% BSA or nonfat dry milk at 4°C overnight. Membranes were incubated with the appropriate horseradish peroxidase–conjugated IgG secondary antibody in TBS-T with 5% nonfat dry milk for 1 hour at room temperature. Immunoreactive protein was visualized using an enhanced chemiluminescence analysis kit (GE HealthCare, Piscataway, NJ).

### Body Composition Analysis

Body composition of mice was determined following sacrifice using whole-body nuclear magnetic resonance imaging (EchoMRI, Echo Medical Systems, Waco, TX).

### Oil-Red-O Staining

Liver samples were immediately frozen after collection. Samples were embedded in O.C.T for sectioning and tissue sections were mounted on glass slides. Oil-Red-O staining was performed on the glass.

### miR-181c Overexpression in Wild-Type Mice

To overexpress miR-181c in WT mice, a miR-181c expression construct was packaged into an AAV-8. Control scrambled miRNA (“Scr”; Cat# AA08-CmiR0001-MR14-100, GeneCopoeia Inc, MD, USA) and miR-181c (“miR-181c OE”; Cat# AA08-MmiR3275-MR14-200, GeneCopoeia Inc, MD, USA) were packaged into a hepatocyte specific serotype, AAV-8 construct. C57BL/6J mice were fed HF diet for 4 weeks prior to being injected with virus construct (10^11^ viral particles in PBS) (n = 8 mice per construct) and remained on HF diet until sacrificed 6 weeks later (Figure 7A). A total of 50 μL of scrambled or miR-181c was injected retro-orbitally. Mice were tested in an ip-GTT 5 weeks after virus injection and sacrificed 1 week later for collection of blood and tissues as described above.

### Statistical Analysis

Data were analyzed by one-way or two-way analysis of variance (ANOVA) and Bonferroni corrected post hoc tests were used where appropriate. All analyses were performed using Prism 9 (GraphPad Software Inc., CA). P<0.05 was considered statistically significant. Results are presented as mean ± standard error of the mean (mean ± SEM).

## Results

### Metabolic profile of mice lacking miR-181c/d

As shown previously, c/d KO mice were born in the expected Mendelian ratios and there were no body weight differences at birth compared to WT littermates [3]. Based on our observed finding that the hearts of c/d KO mice are protected against ischemia-reperfusion injury [3] and diet-induced obesity associated cardiac dysfunction [5], we hypothesized that c/d KO mice would also be protected from metabolic stress in the liver due to HF. Prior to HF exposure, c/d KO mice had increased oxygen consumption (VO_2_; Figure 1A) compared to WT mice (p=0.01), but no differences in CO_2_ production (Figure 1B). This resulted in a lower respiratory exchange ratio (RER), indicating a preferential mobilization of fat, rather than carbohydrates, for energy needs compared to WT mice (Figure 1C, p<0.0001). In addition, c/d KO mice had increased energy expenditure compared to WT mice (Figure 1D, (p=0.01)) on chow diet. However, after 10 and 20 weeks of HFD access, there were no longer any significant group differences in oxygen consumption, CO_2_ production, RER or energy expenditure between the WT and c/d KO mice (Fig. 1A-D). There were significant group x time interactions on VO_2_ (Figure 1A; F (2, 18) = 59.78, p<0.0001), RER (Figure 1C; F (2, 18) = 468.6, p<0.0001), and energy expenditure (Figure 1D; F (2, 18) = 78.54, p<0.0001). There were no significant differences in locomotor activity or food intake between WT and c/d KO mice while on chow or throughout the HF diet exposure period (data not shown).

**Figure 1:**
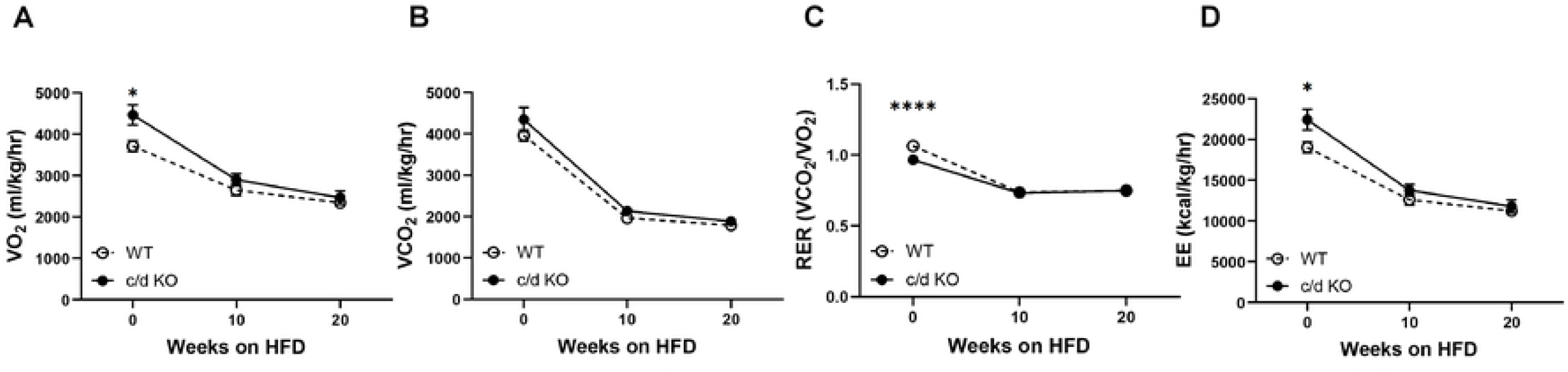
Metabolic profiling of miR-181c/d^-/-^ mice. **(A)** Oxygen consumption (VO_2_), **(B)** carbon dioxide production (VCO_2_), **(C)** respiratory exchange rate (RER), **(D)** energy expenditure (EE) were measured before and during the 26 weeks HF-diet. Data is reflective of the following group sizes: **A-D**, n=4/genotype. Data are expressed as mean ± SEM. Data with differing superscripts differ from each other by p<0.05 by Bonferroni post-hoc analysis after intergroup differences were found by 2-way ANOVA. *p<0.05 or less, two-sample t-test.

### Lack of miR-181c leads to the accumulation of white adipose tissue and impaired glucose tolerance after HFD exposure

While on chow diet (6-8 week of age), there were no differences in body weight between c/d KO and WT mice (Figure 2A). When placed on HFD, both WT and c/d KO mice had elevated body weight compared to their respective chow controls (Figure 2A; F (3, 48) =5.315, p=0.003). After 26 weeks of chow or HF exposure, there was a significant genotype diet effect on body fat (Figure 2B; F (3, 37) = 30.83, P<0.0001). Bonferroni post hoc analysis showed that c/d KO-HF mice had increased body fat compared to WT-CH, c/d KO-CH and WT-HF mice. In addition, there were significant effects of genotype and diet with respect to lean mass (Figure 2C; F (3, 40) = 28.44, p<0.0001). c/d KO-CH mice had lower lean body mass compared to WT-CH mice, but not WT-HF mice. When exposed to HF for 26 weeks, c/d KO-HF mice had lower lean mass as a percent of total carcass weight compared to all groups.

**Figure 2:**
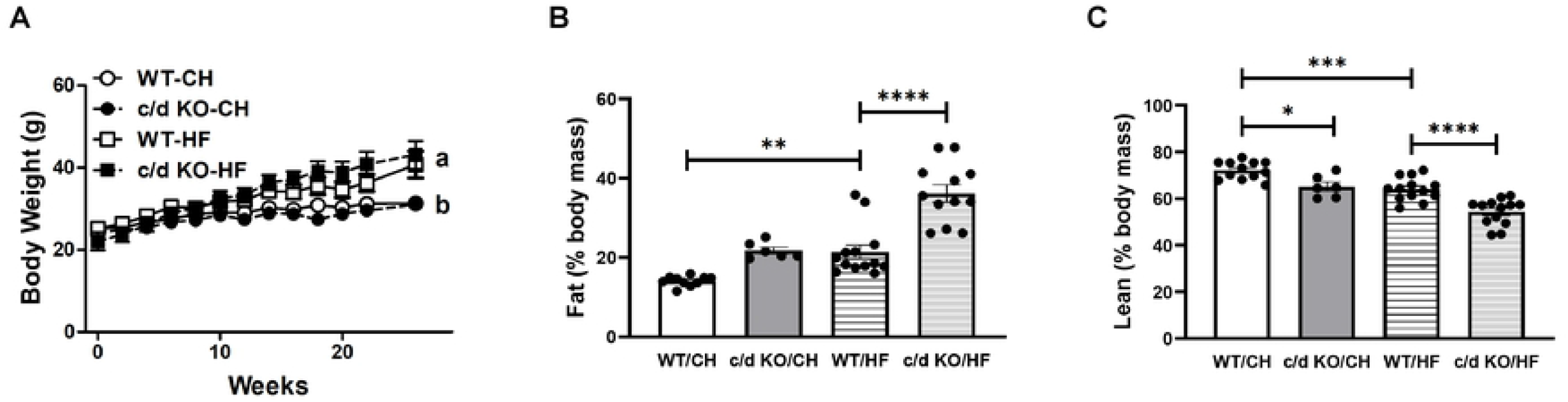
Effect of High Fat (HF)-diet on miR-181c/d^-/-^ mice. **(A)** body weight, **(B)** percent fat mass, **(C)** percent lean mass, were measured before and during the 26 weeks HF-diet. Data is reflective of the following group sizes: **A**, n=4/genotype; **B-C**, n=8-13/genotype/diet. Data are expressed as mean ± SEM. Data with differing superscripts differ from each other by p<0.05 by Bonferroni post-hoc analysis after intergroup differences were found by 2-way ANOVA. *p<0.05 or less, two-sample t-test.

At baseline, there were no significant differences in blood glucose levels between WT and c/d KO mice (Figure 3A). However, baseline plasma insulin levels were significantly higher in the c/d KO mice compared to WT (p=0.03) (Figure 3B). When challenged with a glucose tolerance test, there was an overall time by group interaction effect on plasma glucose levels (Figure 3A; F (5, 72) = 24.21, p<0.0001). Bonferroni post-hoc correction analysis showed that at 45-, 60- and 120-minutes post-glucose injection, circulating blood glucose levels were higher in c/d KO mice compared to WT mice. In addition, c/d KO mice had an elevated AUC (p=0.004) compared to WT mice. There was a main effect of group (F (1, 71) = 18.47, p<0.0001) and time (F (5, 71) = 2.714, p=0.0266) for plasma insulin levels (Figure 3B). There were no statistically significant differences in the AUC between WT and c/d KO mice.

**Figure 3:**
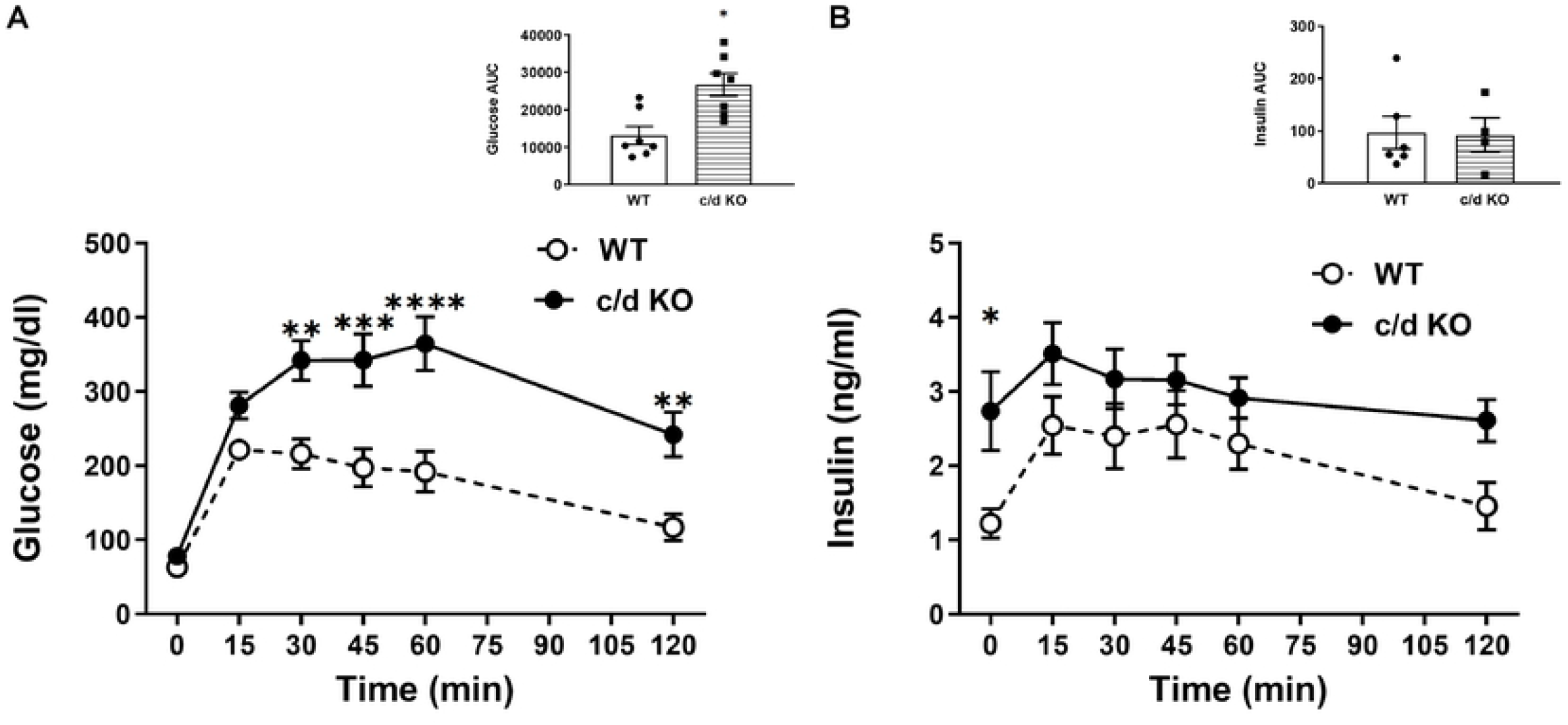
Metabolic consequences of HF-diet on miR-181c/d^-/-^ mice. **(A)** blood glucose and area under the curve during an ip-gluocse tolerance test (GTT) and (**B)** plasma insulin levels and area under the curve during an ip-GTT. Data is reflective of the following group sizes: n=6-7/genotype/diet. Data are expressed as mean ± SEM. Data with differing superscripts differ from each other by p<0.05 by Bonferroni post-hoc analysis after intergroup differences were found by 2-way ANOVA. *p<0.05 or less, two-sample t-test.

### Lack of miR-181c leads to impaired lipid biosynthesis in the liver after HFD exposure

After 26 weeks of HF diet exposure, we observed elevated lipid accumulation in the liver of c/d KO compared to WT mice as measured by oil-red-o staining (Figure 4A). There were no differences in plasma leptin levels between c/d KO and WT mice while on chow diet (Figure 4B). However, after exposure to HF diet, both WT and c/d KO mice had elevated plasma leptin levels (F (3, 19) = 53.43, p<0.0001). There was a main effect of diet (F (3, 18) = 19.79, p<0.0001) on plasma triglyceride levels (Figure 4C). Post hoc analysis showed that HF fed c/d KO mice had elevated plasma triglyceride levels compared to chow-fed WT mice. Post hoc analysis revealed that within the HFD groups, liver triglyceride levels were significantly higher in the c/d KO compared to WT mice and respective chow-fed controls.

**Figure 4:**
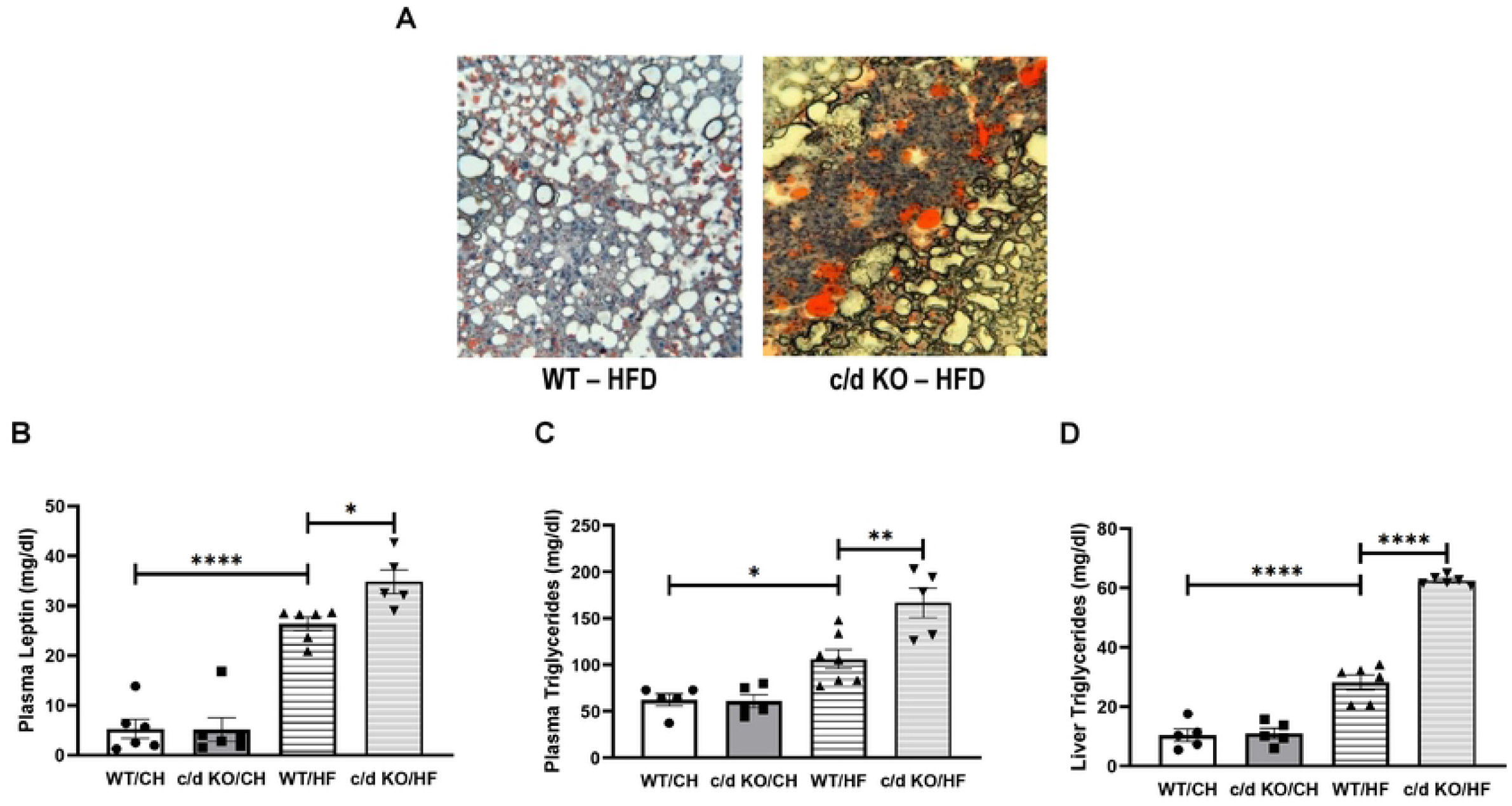
Effect of High Fat (HF)-diet on the liver of miR-181c/d^-/-^ mice. **(A)** liver oil-red-o staining for lipid droplets, **(B)** plasma leptin levels, **(C)** plasma triglyceride levels, and **(D)** liver triglyceride content were analyzed from the normal chow and HF-fed mice from WT and c/d KO groups. Data is reflective of the following group sizes: n=5-7/genotype/diet. Data are expressed as mean ± SEM. Data with differing superscripts differ from each other by p<0.05 by Bonferroni post-hoc analysis after intergroup differences were found by 2-way ANOVA. *p<0.05 or less, two-sample t-test.

### Effect of HF diet on liver miR-181c expression

Recent data shows that the HF upregulates miR-181c in the heart [5]. To determine the regulation of miR-181c in the liver during diet-induced metabolic stress, we measured the expression of miR181c in WT-mice after exposure to 26 weeks of HF diet. Interestingly, miR-181c was decreased in the liver of WT mice after exposure to HF (Figure 5A; p=0.02). In the heart, miR-181c mainly translocate to the mitochondrial fraction [1–3]. Whole tissue analysis in WT mice showed the expression of miR-181c in the liver was markedly higher than heart tissue (Figure 5B). Next, to detect miR-181c in liver mitochondria, we isolated mitochondrial miRNA-enriched total RNA from both the heart and liver mitochondrial fraction. We detected significantly higher levels of miR-181c expression in the mouse heart mitochondria, compared to liver mitochondria (Figure 5C). Therefore, taken together these data suggest that unlike heart, miR-181c is mainly expressed in the cytosolic fraction in the liver.

**Figure 5:**
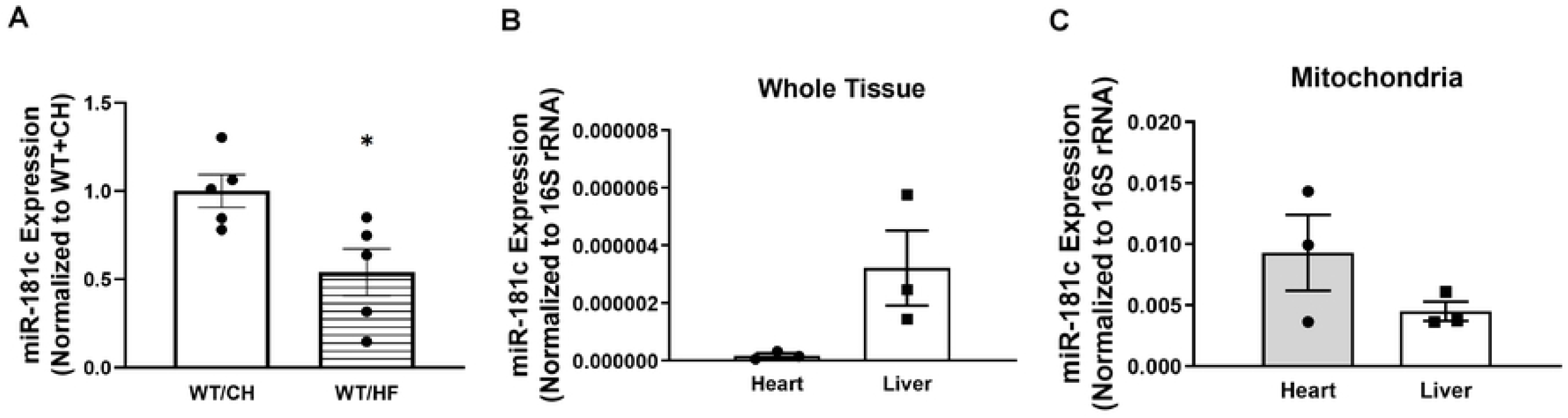
Cytosolic localization of miR-181c in the Liver. Quantitative polymerase chain reaction (qPCR) (SYBR kit) analysis of miR-181c expression **(A)** in total RNA from the liver of normal chow (CH)-fed and high fat (HF)-fed WT mice, **(B)** in total RNA from the heart and liver tissue, and **(C)** the mitochondrial pellets of WT heart and liver tissue. Data is reflective of the following group sizes: n=3-7/genotype/diet. Data are expressed as mean ± SEM. Data with differing superscripts differ from each other by p<0.05 by Bonferroni post-hoc analysis after intergroup differences were found by 2-way ANOVA. *p<0.05 or less, two-sample t-test.

### miR-181c targets IDH 1 in the cytoplasm to inhibit lipid biosynthesis in the liver

Previously, it was demonstrated that the “seed” sequence for miR-181 can directly bind to the 3’-UTR of the isocitrate dehydrogenase 1 (IDH 1) gene at one putative site (UGAAUGU) [29]. In addition, overexpression of cytosolic-specific IDH (IDHc) resulted in elevated liver triglycerides and adipose tissue accumulation [30]. Therefore, since miR-181c is mainly expressed in the cytosolic fraction of the liver (Figures 5B and 5C) and IDHc is involved in lipid biosynthesis in the liver, we measured IDH protein expression in the liver of chow-fed WT and c/d KO mice. The expression of IDH 1 is in the cytoplasm; on the other hand, IDH 2 is expressed in the mitochondrial fraction. Therefore, we measured the protein expressions of both IDH 1 and IDH 2 in c/d KO mouse liver and compared with WT liver. Loss of miR-181c results in the up-regulation of IDH 1 protein in the liver (Figure 6A) compared to WT mice (p=0.015). However, there were no changes in IDH 2 protein expression after loss of miR181c (Figure 6B) compared to WT mice. This data suggests that miR-181c targets IDH 1 mRNA in the cytoplasmic fraction of the hepatocytes.

**Figure 6:**
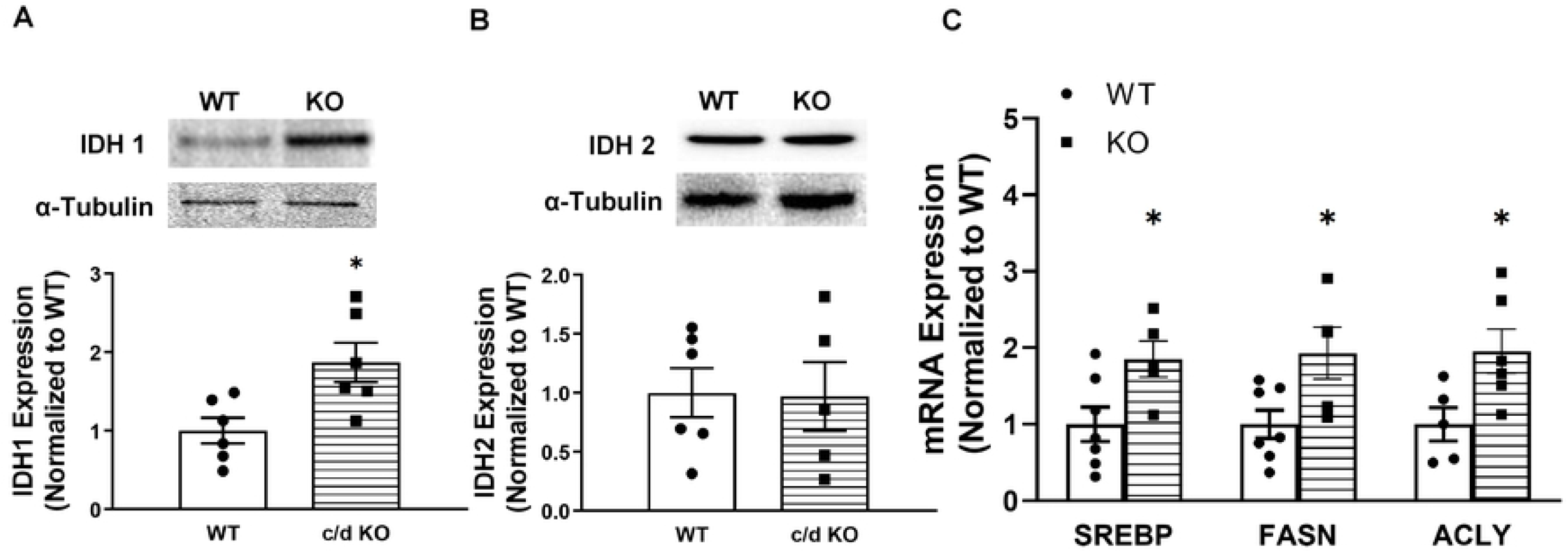
Lack of miR-181c activate lipogenesis by upregulating IDH 1. **(A)** IDH 1 protein expression and **(B)** IDH 2 protein expression in the liver. Quantitative polymerase chain reaction (qPCR) of liver mRNA expression of genes involved in lipogenesis. **(C)** Sterol regulatory element binding transcription factor 1 (SREBP1), fatty acid synthase (FASN) and ATP citrate lyase (ACLY). Data is reflective of the following group sizes: n=5-7/genotype/diet. Data are expressed as mean ± SEM. Data with differing superscripts differ from each other by p<0.05 by Bonferroni post-hoc analysis after intergroup differences were found by 2-way ANOVA. *p<0.05 or less, two-sample t-test.

Finally, to determine potential mechanisms, which may contribute to impaired metabolic functioning in the liver when placed on HF diet via IDH 1 pathway, we measured liver mRNA expression of genes involved in lipogenesis in chow-fed WT and c/d KO mice (Figure 6C). Liver mRNA expression of genes involved in de novo lipid synthesis (ACLY; p=0.03, FASN; p=0.03) and fatty acid synthesis (SREBP1; p=0.03) were increased c/d KO mice compared to WT controls.

### Liver-specific miR-181c overexpression

Next, to determine if overexpression of miR-181c can protect against the liver-specific metabolic consequences due to HF exposure, we used an AAV (serotype 8) package system. First, the dose of viral particles/retro-orbital vein delivery and optimal time points were optimized (Figure 7A). After 4 weeks of HF diet, the mice were randomly selected for either AAV-8 scramble or AAV-8-miR-181c injection. Both groups of mice then kept on HFD for additional 6 weeks. Using this treatment regimen of *in vivo* miR181c delivery, we successfully overexpressed miR-181c in the liver by 2.5-fold (Figure 7B, p=0.03). Importantly, there were no overexpression of miR-181c in heart (Figure 7C), lung (Figure 7D), spleen (Figure 7E) and kidney (Figure 7F) with AAV-8-miR-181c injected group compared to AAV-8 scramble injected group.

**Figure 7:**
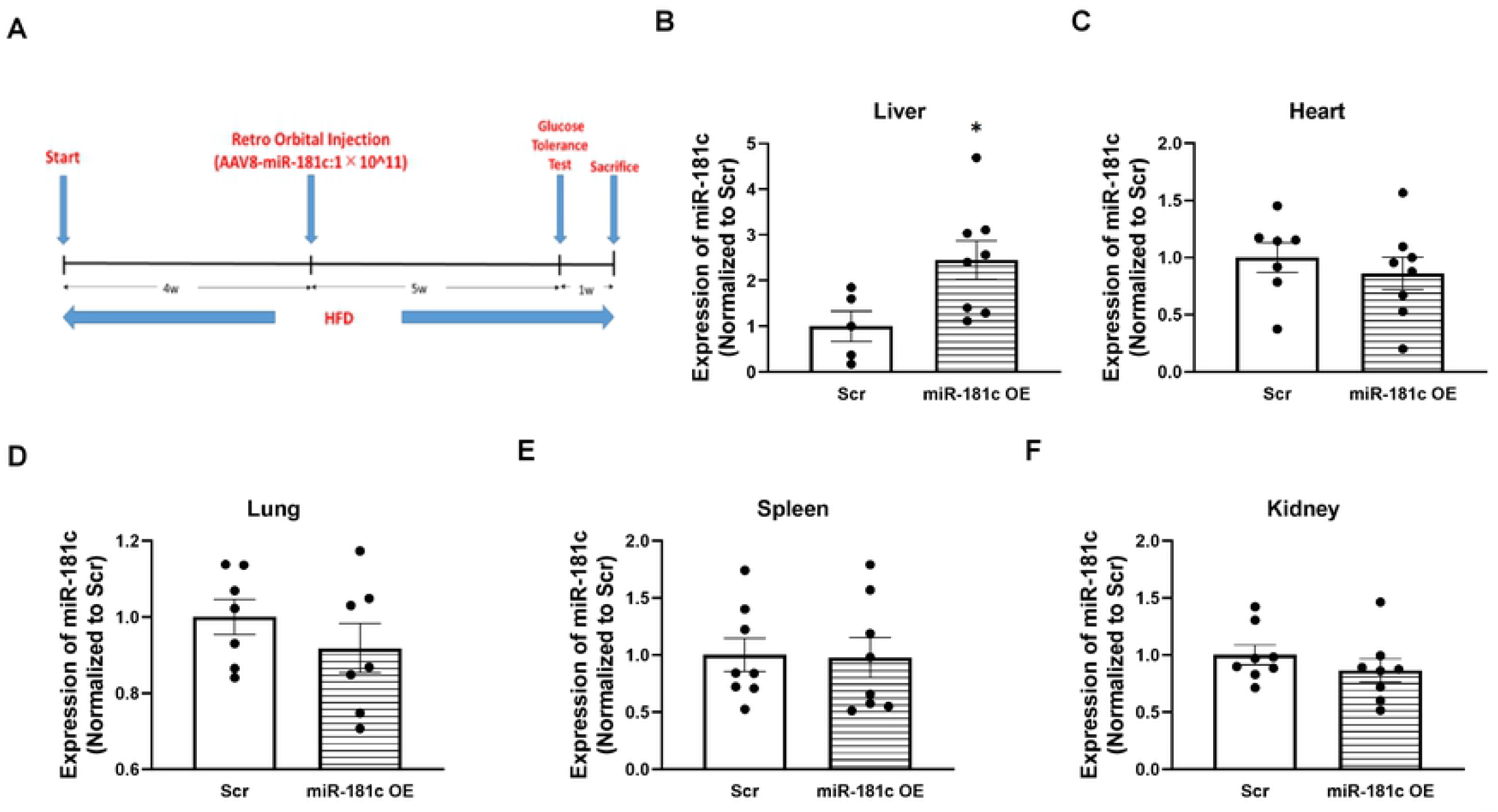
Liver-specific miR-181c delivery. profiling of miR-181c/d^-/-^ mice. **(A)** *in vivo* miR-181c overexpression protocol. Quantitative polymerase chain reaction (qPCR) (SYBR kit) analysis of miR-181c expression in total RNA isolated from **(B)** liver, **(C)** heart, **(D)** lung, **(E)** spleen and **(F)** kidney. Data is reflective of the following group sizes: **B-F** n=8. Data are expressed as mean ± SEM. Data with differing superscripts differ from each other by p<0.05 by Bonferroni post-hoc analysis after intergroup differences were found by 2-way ANOVA. *p<0.05 or less, two-sample t-test.

There was no body weight difference between AAV-8-miR-181c and AAV-8 scramble injected groups after 10 weeks of HF diet (Figure 8A). Furthermore, at baseline, there were no significant differences in blood glucose (Figure 8B) and insulin (Figure 8C) levels between AAV-8-miR-181c and AAV-8 scramble groups. When challenged with a glucose tolerance test, there was an overall time by group interaction effect on plasma glucose level (Figure 8B). There were no circulating blood glucose or insulin levels between AAV-8-miR-181c and AAV-8 scramble groups.

**Figure 8:**
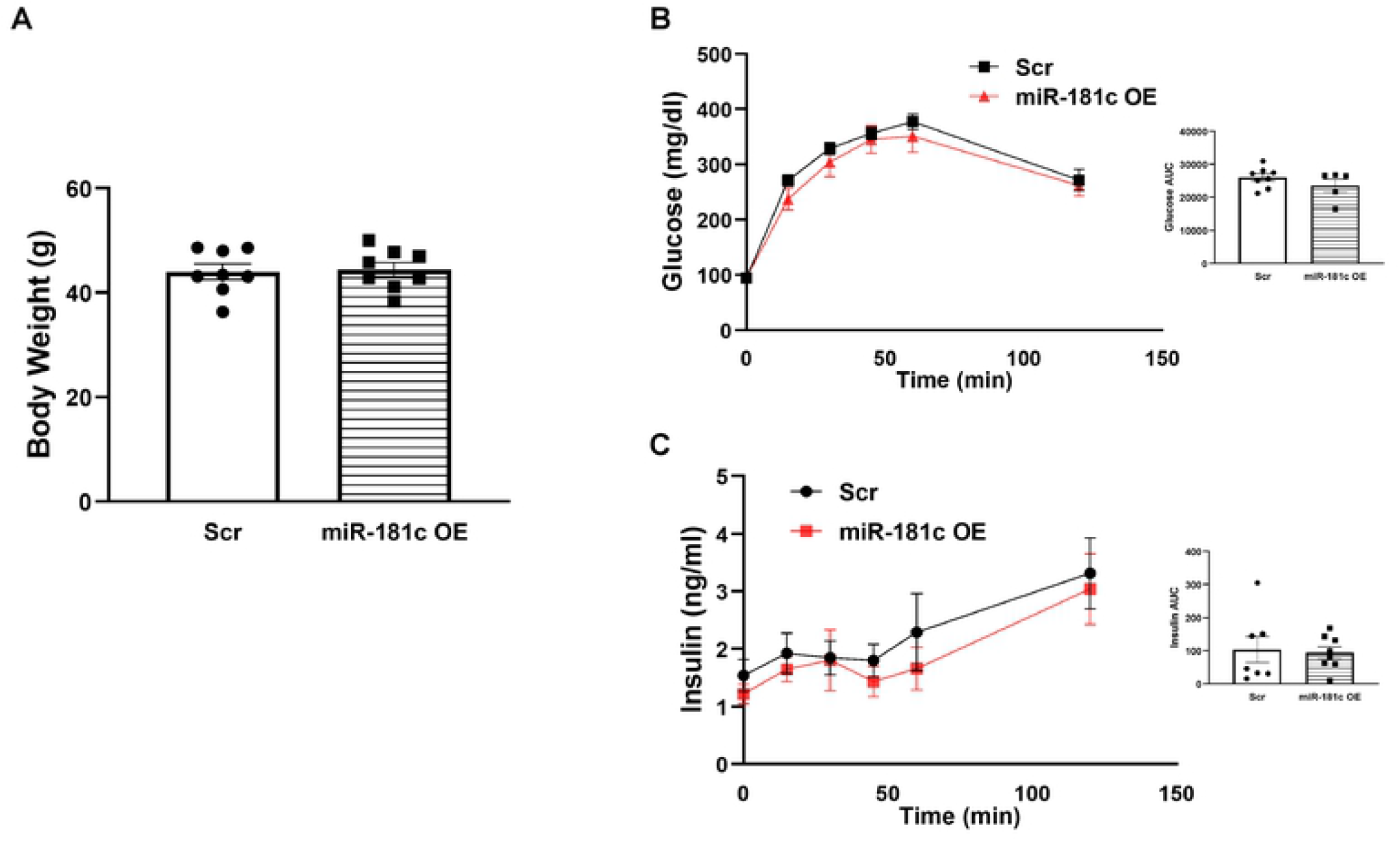
Effect on insulin secretion by miR-181c overexpression in the liver. **(A)** body weight, (**B)** blood glucose and area under the curve during an ip-GTT, and **(C)** plasma insulin levels and area under the curve during an ip-GTT. Data is reflective of the following group sizes: n=8. Data are expressed as mean ± SEM. Data with differing superscripts differ from each other by p<0.05 by Bonferroni post-hoc analysis after intergroup differences were found by 2-way ANOVA. *p<0.05 or less, two-sample t-test.

### Overexpression of miR-181c mitigates the lipogenesis from the HFD exposure

Liver-specific overexpression of miR-181c significantly downregulates IDH 1 expression (Figure 9A) without altering IDH 2 expression (Figure 9B), after 10 weeks of HF diet. Given our findings of the potential role for miR-181c/d in IDH 1 regulated lipogenesis, we measured the expression of genes involved in lipogenesis in the liver. After 10 weeks of HFD, treatment with miR-181c resulted in lower liver mRNA expression of genes involved in long chain fatty acid synthesis (FASN; p=0.03, SREBP1; p=0.003) (Figure 9C).

**Figure 9:**
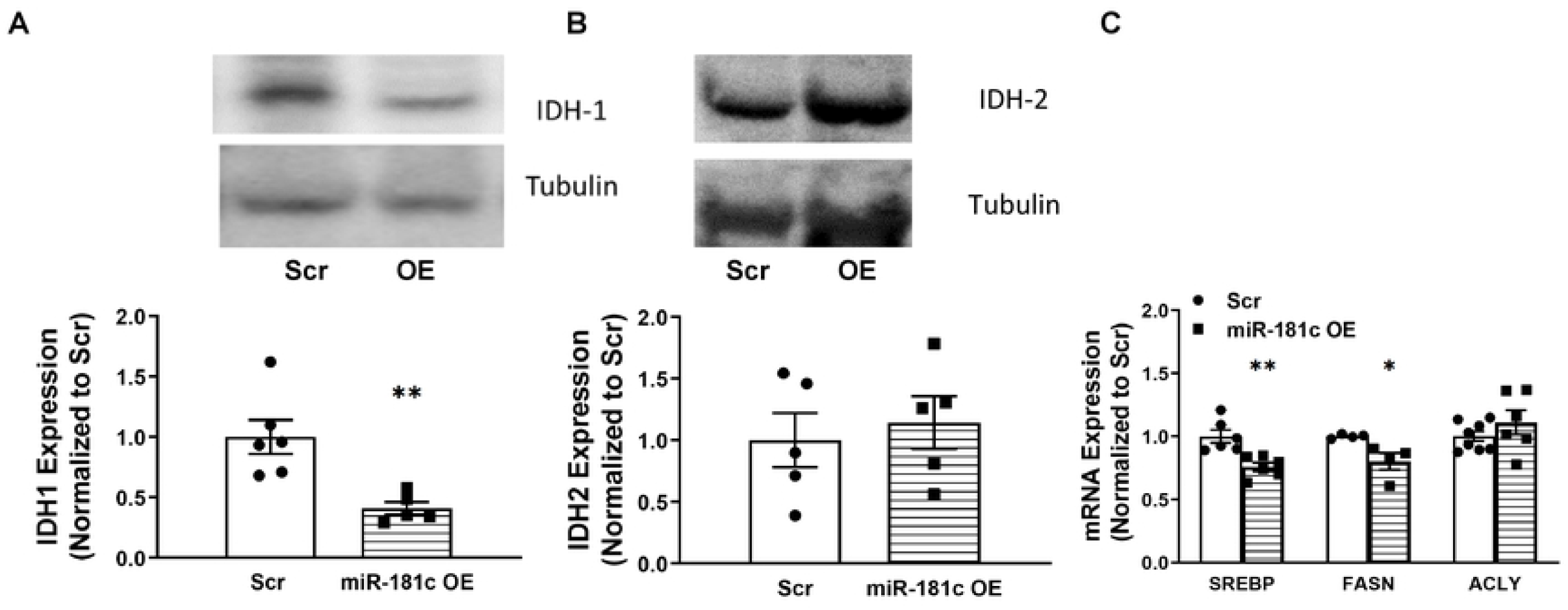
Effect of liver-specific miR-181c overexpression on lipogenesis. **(A)** IDH 1 protein expression and **(B)** IDH 2 protein expression in the liver. Quantitative polymerase chain reaction (qPCR) of liver mRNA expression of genes involved in lipogenesis. **(C)** Sterol regulatory element binding transcription factor 1 (SREBP1), fatty acid synthase (FASN) and ATP citrate lyase (ACLY). Data is reflective of the following group sizes: n=8. Data are expressed as mean ± SEM. Data with differing superscripts differ from each other by p<0.05 by Bonferroni post-hoc analysis after intergroup differences were found by 2-way ANOVA. *p<0.05 or less, two-sample t-test.

After 10 weeks of HF exposure, we observed elevated lipid accumulation in the liver of AAV-8 scramble injected mice by oil-red-o staining (Figure 10A, left panel); however, AAV-8-miR-181c injected mice shows a significantly lower lipid droplet (Figure 10A, right panel). Post hoc analysis revealed that within the HFD groups, liver triglyceride levels were significantly lower in the AAV-8-miR-181c injected mice compared to AAV-8 scramble injected mice (Figure 10B). Finally, among the HF diet group liver-specific overexpression of miR-181c can also lower the plasma triglyceride (Figure 10C) and plasma leptin (Figure 10D) levels compared to AAV-8 scramble injected mice.

**Figure 10:**
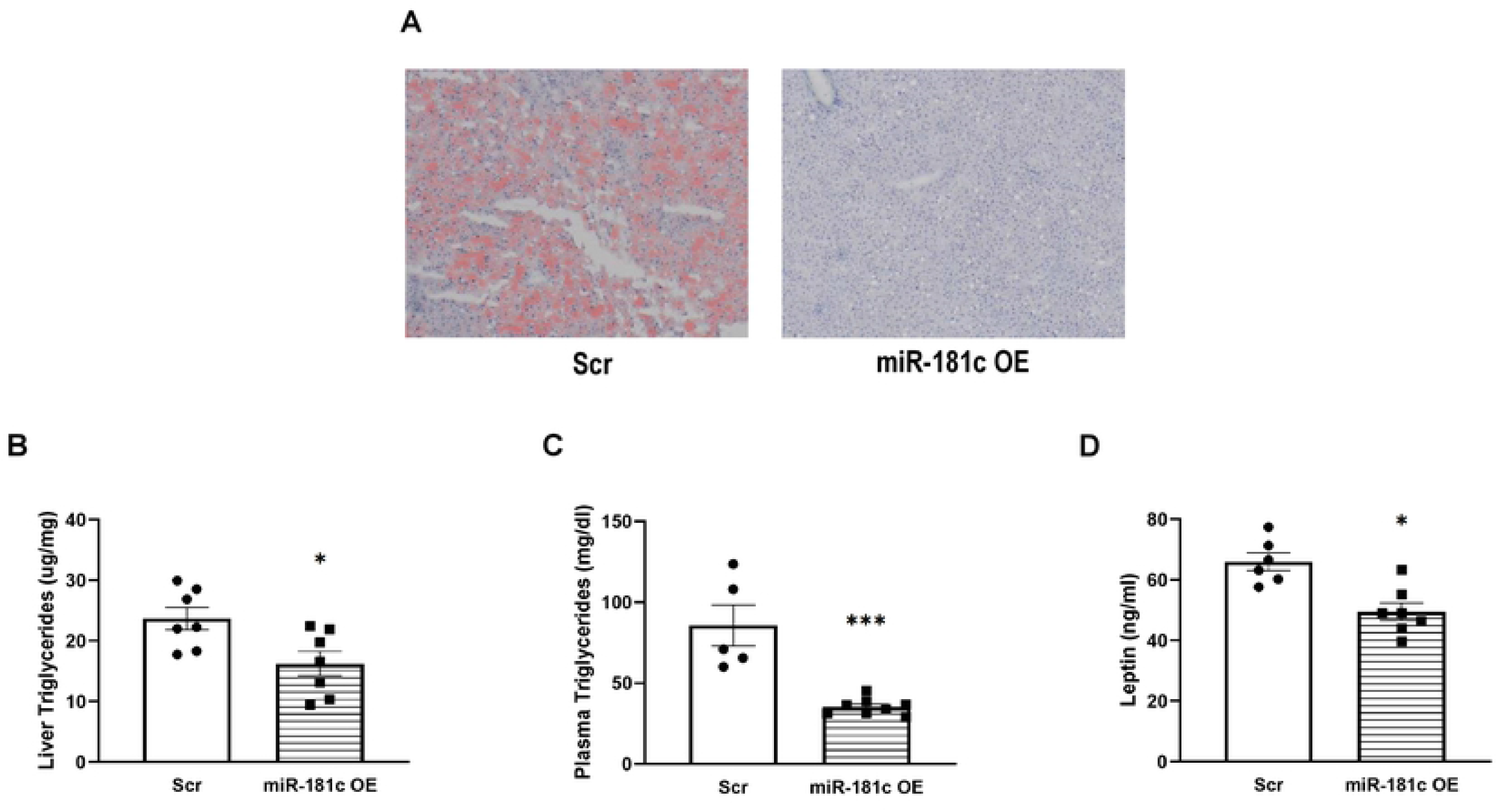
Effect of liver-specific miR-181c overexpression on metabolic consequences of a high-fat diet. **(A)** liver oil-red-o staining for lipid droplets, **(B)** liver triglyceride content, **(C)** plasma triglyceride levels, and **(D)** plasma leptin levels were analyzed from the scramble and miR-181c overexpressed WT mice of 10-week high fat diet. Data is reflective of the following group sizes: n=8. Data are expressed as mean ± SEM. Data with differing superscripts differ from each other by p<0.05 by Bonferroni post-hoc analysis after intergroup differences were found by 2-way ANOVA. *p<0.05 or less, two-sample t-test.

## Discussion

The ability of miR-181c overexpression to confer robust protection in the obesity paradigm has two important implications. First, these results support miR-181c as a potential therapeutic target to combat obesity and lipogenesis. As it has already been established that miR-181c can be delivered *in vivo* using a nanovector [1], these findings identify a paradigm useful for testing the efficacy of miR-181c overexpression and determine its role in fat inhibition and cholesterol biosynthesis. Second, these findings demonstrate that compensation for the loss of miR-181c can confer protection against obesity when exposed to a HFD. miR-181c/d^-/-^ mice have a severe obese phenotype when fed HFD. The data from the miR-181c/d^-/-^ mice with HFD and *in vivo* liver-specific overexpression of miR-181c exposed to HFD, suggest miR-181c can offer protection in the liver from HFD exposure by inhibiting lipogenesis.

The nuclear-encoded miR-181 family members play an important role in cardiac function by regulating target genes in both the cytoplasm and within the mitochondria. miR-181a/b regulates PTEN expression in the cytoplasm, whereas miR-181c regulates the mitochondrial gene mt-COX1 in the mitochondria [3]. We have previously demonstrated, both *in vitro* [2] and *in vivo* [1], a pivotal role for mitochondrial miR-181c in cardiac dysfunction. We [1–3] and others [4], have identified a significant role for miR-181 in the heart during end-stage heart failure. In the present study, we identified a nuclear-encoded cytoplasmic target, IDH 1, for miR-181c in the liver (another mitochondria-enriched organ). Mature miR-181c can be expressed mainly in the mitochondrial fraction of cardiomyocytes; however, in hepatocytes, miR-181c directly binds to the 3’-UTR of IDH 1 in the cytoplasm. By down-regulating IDH 1 during DIO, miR-181c can regulate lipid metabolism.

Acetyl-CoA and NADPH are essential common precursors and cofactors for lipid biosynthesis. IDH 1, a key NADPH producer, contributes to the activation of triglyceride and cholesterol synthesis by the liver [30]. Similar to the metabolic phenotype of IDH 1 Tg mice [30], miR-181c/d^-/-^ mice have, an increase in body fat composition and insulin insensitivity after 26 weeks of a HFD. These two mouse models highlight the potential role of IDH 1 in the induction of fatty liver, hyperlipidemia, and obesity by alteration of lipid biosynthesis in the liver, especially under HFD-induced stress. Furthermore, IDH 1 has also been demonstrated to activate lipogenesis under hypoxic conditions by reductive glutamine metabolism [31]. IDH 1 can also regulate the lipogenic pathways by activating the transcriptional factors SREBP1 and SREBP2 [32]. Alternatively, Chu et al. [29], showed that knockdown of IDH 1 downregulates lipid synthesis-related gene expression and up-regulates β-oxidation and cholesterol transport-related gene expression, thereby inhibiting lipid synthesis in the liver. Therefore, liver-specific overexpression of miR-181c may be a potential therapeutic target for abnormal fat synthesis during DIO. miR-181c can attenuates lipid biosynthesis by directly bind to the 3’UTR of IDH 1 mRNA and perhaps miR-181c may play that therapeutic molecule (Figure 11).

**Figure 11:**
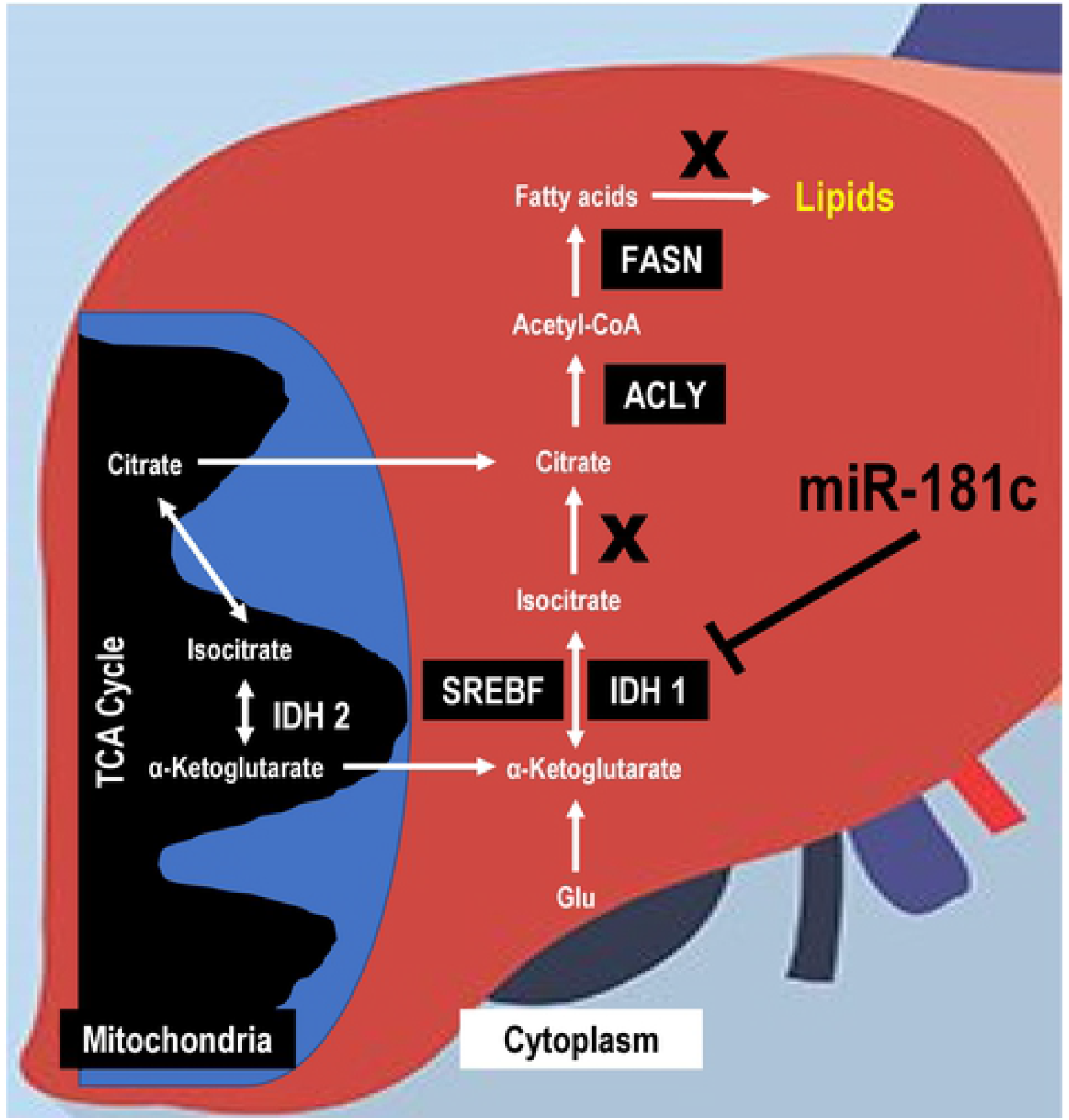
Role of miR-181c in regulating lipogenesis by high fat diet. This schematic diagram illustrates key steps in the signaling pathway linking high fat diet with increased lipogenesis. According to this model, high fat triggers lipogenesis by activating IDH 1 though SREBF, ACLY and FASN pathway. Liver-specific miR-181c can directly binds to 3’UTR of IDH 1 mRNA and mitigate lipogenesis.

In this study, we have observed that miR-181c/d^-/-^ mice show a severe obese phenotype with significant hyperinsulinemia and hyperglycemia once we switched to the 60% HFD. These data also suggest that the miR-181c-IDH 1 pathway plays an important role in dietary-stress. Thus, liver-specific overexpression of miR-181c can protect from dietary-stress by inhibiting lipogenesis. miRNA therapeutics may be a potential treatment regimen, however, the tissue-targeted delivery is challenging as there is potential for off-target side effects [33]. In the obesity field, miR-143~145 cluster delivery in the liver has been shown to have a protective effect against obesity-associated diabetes [34]. Previously, we have demonstrated that miR-181c can be overexpressed *in vivo*, using an electrostatic complex: nanovector with positively charged liposomal nanoparticles and negatively charged plasmid DNA. Previous work also shows no immune response during this nanovector treatment regimen [1]. However, nanovector-based miR-181c delivery can deliver miR-181c in all the tissues, including heart. Overexpression of miR-181c can cause severe cardiac dysfunction via mitochondrial mechanisms [1–3]. In this study, we packaged miR-181c construct in AAV-8 and injected 10^11^ viral particle/mouse through retro-orbital injection, which has failed to overexpress miR-181c in heart, lung, spleen and kidney (Figure 7). Thus, the future technologies should focus on liver-specific miR-181c delivery to combat against DIO complications, such as nonalcoholic fatty liver disease. The future of such intervention might be useful for the patients with nonalcoholic fatty liver disease.

## Conclusions

In summary, in this present study we have demonstrated that miR-181c plays an essential role in inhibiting triglyceride and cholesterol biosynthesis by targeting IDH 1 mRNA in the cytoplasm. miR-181c/d^-/-^ mice have a severe obese phenotype during HFD. Overexpression of miR-181c during DIO can protect from the metabolic consequences of HFD exposure by altering lipid metabolism.

## Acknowledgments

This work was supported by grants from the MSCRF, Mscrfd-4313, U54AG062333 and U18TR003780 (S.D.), 5R01HL039752 (C.S.), JHU-UMD Diabetes Research and Training Center (NIDDK P30 DK079637).

## Non-Standard Abbreviations and Acronyms

miRNA: microRNA
miR-181c/d^-/-^ mice: miR-181c and d knock-out mice
IDH 1: Isocitrate Dehydrogenase 1
HFD: 60% High fat diet
DIO: Diet-Induced Obesity
ABCA1: ATP-binding cassette, subfamily A, member 1
ABCG1: ATP-binding cassette, subfamily G, member 1
ACLY: ATP citrate lyase
APOE: Apolipoprotein E
CPT1A: Carnitine Palmitoyltransferase 1A
FASN: Fatty Acid Synthase
GAPDH: Glyceraldehyde-3-Phosphate Dehydrogenase
HADHB: Hydroxyacyl-Coenzyme A Dehydrogenase Trifunctional Multienzyme Complex, β subunit
SREBP1: Sterol Regulatory Element Binding Transcription Factor 1
SNORD6: Small Nucleolar RNA, C/D Box 6

